# Impaired BCAA catabolism during adipocyte differentiation decreases glycolytic flux

**DOI:** 10.1101/2022.05.27.493780

**Authors:** Courtney R. Green, Karl A. Wessendorf-Rodriguez, Rory Turner, Justin D. Hover, Anne N. Murphy, Christian M. Metallo, Martina Wallace

## Abstract

Dysregulated branched chain amino acid (BCAA) metabolism has emerged as a key metabolic feature associated with the obese insulin resistant state, and adipose BCAA catabolism is decreased in this context. BCAA catabolism is upregulated early in adipogenesis, but the impact of suppressing this pathway on the broader metabolic functions of the resultant adipocyte remain unclear. Here, we use CRISPR/Cas9 to target *Bckdha* and *Acad8* in pre-adipocytes and induce a deficiency in BCAA or valine catabolism through differentiation. We characterise the transcriptional and metabolic phenotype of these cells using RNAseq and ^13^C metabolic flux analysis within a network spanning glycolysis, tricarboxylic (TCA) acid metabolism, BCAA catabolism, and fatty acid synthesis. While lipid droplet accumulation is maintained in *Bckdha*-deficient adipocytes, they display a more fibroblast-like transcriptional signature. In contrast, *Acad8* deficiency minimally impacts gene expression. Decreased glycolytic flux emerges as the most distinct metabolic feature of *Bckdha*-deficient cells, accompanied by a ∼40% decrease in lactate secretion, yet pyruvate oxidation and utilization for *de novo* lipogenesis are increased to compensate for loss of BCAA carbon. Glutamine anaplerosis was also increased, though we observed a general decrease in levels of most non-essential amino acids consistent with an impact on nitrogen homeostasis. Overall, our data suggest that both metabolic and regulatory cross-talk exists between BCAA catabolism, glycolysis, and nitrogen metabolism in differentiated adipocytes. Suppression of BCAA catabolism associated with metabolic syndrome may result in a metabolically compromised adipocyte.

## Introduction

The branched-chain amino acids (BCAAs) leucine, isoleucine, and valine are essential dietary amino acids whose elevated circulating levels are emerging as a hallmark of metabolic syndrome (1–3). It has become evident that BCAA metabolism is dysregulated in multiple different tissues in this context and this includes adipose tissue where transcriptional and proteomic studies in humans have shown that BCAA catabolism is significantly suppressed with obesity and insulin resistance (4, 5). In addition, in cross tissue comparisons of BCAA catabolic flux in lean and obese mice, adipose tissue displays one of the biggest decreases in BCAA breakdown (6, 7). The BCAA pathway is one of the most consistently altered metabolic pathways in the context of adipose dysfunction and thus it is important to develop our knowledge of the role this pathway plays in adipose tissue.

BCAA catabolism is upregulated during adipocyte differentiation (4, 8–14) and it is initiated by mitochondrial BCAT2, which transfers the amino nitrogen to α-ketoglutarate yielding glutamate and a corresponding branched-chain α-keto acid (BCKA). These BCKAs are subsequently oxidized by the branched-chain ketoacid dehydrogenase (BCKD) complex, a highly regulated, rate-limiting step in BCAA catabolism. A series of enzymatic steps further catabolize short branched-chain fatty acid (SBCFA)-CoAs to acetyl-CoA and Succinyl-CoA which can enter the TCA cycle. In addition to serving as a source of nitrogen and carbon for the cell, many intermediates of BCAA catabolism have been proposed to play signaling roles or serve as post-translational modifications and thus BCAA catabolism may impact cell function in multiple ways (7, 15–18).

We and others have previously investigated how genetically altering BCAA metabolism impacts adipocyte differentiation and found that shRNA mediated knockdown of *Bcat2* or *Bckdha* (14), the leucine catabolic enzymes *Mccc1* (11), and siRNA-mediated knockdown of the valine catabolic enzyme *Hibch* (19) throughout differentiation decreased lipid accumulation indicating impaired differentiation. In mice, disruption of BCAA catabolism in white adipocytes via knockout has yielded varying results in overall phenotype but enhanced BCAA catabolism has generally been associated with increased white adipose depot size in response to high fat diet (16, 20).

Collectively, these data indicate altered BCAA catabolism impacts adipogenesis and adipose mass but a comprehensive assessment of how this impacts broader metabolism in adipocytes is lacking. Here we use CRISPR/Cas9 to generate pre-adipocytes deficient in *Bckdha* or *Acad8,* followed by metabolic characterization of resultant adipocytes using RNAseq, stable isotope tracing and metabolic flux analysis. We find that glucose uptake is substantially decreased indicating crosstalk between BCAA catabolism and signaling mechanisms necessary to upregulate glycolysis during differentiation. However, *de novo* lipogenesis is maintained via enhanced pyruvate utilisation at the expense of lactate production. This study demonstrates how BCAA catabolism during adipogenesis is required for the generation of metabolically competent adipocytes.

## Materials & Methods

### Cell Culture & Differentiation

All reagents were purchased from Sigma-Aldrich unless otherwise noted. All media and sera were purchased from Life Technologies unless otherwise stated. Mouse 3T3-L1 pre-adipocytes were provided by Prof. Alan Saltiel and cultured in high glucose Dulbecco’s modified Eagle medium (DMEM) (Life Technologies) supplemented with 10% bovine calf serum (BCS) below 70% confluence. Cells were regularly screened for mycoplasma contamination. For differentiation, 10,000 cells/well were seeded onto 12-well plates and allowed to reach confluence (termed Day -1). On Day 0, differentiation was induced with 0.5 mM 3-isobutyl-1-methylxanthine (IBMX), 0.25 μM dexamethasone, and 1 μg/ml insulin in DMEM containing 10% FBS. Medium was changed on Day 3 to DMEM + 10% FBS with 1 μg/ml insulin. Day 6, and thereafter, DMEM + 10% FBS was used.

Isotope tracing was carried out 7d post-induction of differentiation. Cells were incubated in custom DMEM (Hyclone) in which the metabolite specified was replaced with the ^13^C- or ^2^H-labeled version for 24h unless otherwise specified. Fatty acids were then either calculated as percent total fatty acids and normalized to control conditions or normalized to [^2^H_31_]Palmitate internal standard and normalized to control conditions.

### Generation of lentiviral CRISPR/Cas9 KO 3T3-L1 adipocytes

Control, Bckdha, and Acad8 target sequences (Supplementary Table 4) were cloned into the lentiCRISPRv2 plasmid, a gift from Feng Zhang (Addgene plasmid #52961) (21). For lentivirus production, 2-2.5 million HEK293FT cells were placed in 10cm tissue culture plates in 2 mL of DMEM (containing 1% penicillin/streptomycin, 10% FBS). 24h later, transfection was performed using Lipofectamine 3000 (Invitrogen) with 1.3μg VSV.G/PMD2.G, 5.3μg of lenti-gag/pol/PCMVR8.2 and 4μg of lentiviral vector. Lentivirus-containing supernatants were harvested 48 and 72h later, combined, and concentrated using Amicon Ultra-15 centrifugal filters, 100,000 NMWL (Millipore) following the manufacturer’s protocol. 3T3-L1 pre-adipocytes were infected with 7μL of virus in 2mL of medium containing 7.5μg of polybrene in a 35mm cell culture dish. Media was changed the following day and allowed to recover for 24h. Selection was performed using 2μg/mL puromycin. Cells were then plated to 12-well plates for differentiation as described above. Puromycin was removed from the medium beginning on day 0 of differentiation.

### Insulin-stimulated glucose metabolism

Differentiated 3T3-L1 adipocytes were insulin-starved for 6h in 5mM glucose DMEM + 0.5% FBS. Cells were then glucose-starved in Kreb’s Ringer Buffer (KRB) + 1.26g/L sodium bicarbonate for 1h. Cells were labeled using 0.5mL 5mM [U-^13^C_6_]glucose KRB ± 100nM insulin for 10 minutes before extraction for GC-MS analysis.

### Western Blots

3T3-L1 adipocytes were lysed in ice-cold RIPA buffer with 1x protease inhibitor (Sigma-Aldrich). 25μg total protein was separated on a 10% SDS-PAGE gel for BCKDHA and β-actin, while 50μg total protein was loaded for ACAD8. The proteins were transferred to a nitrocellulose membrane and immunoblotted with anti-BCKDHA (Novus Biologicals NBP1-79616) (1:2,500 dilution), anti-β-Actin (Cell Signaling 4970S) (1:10,000), anti-ACAD8 (Aviva Systems Biology OAAB06085) (1:1,000 dilution). Specific signal was detected with horseradish peroxidase-conjugated secondary antibody goat anti-rabbit (1:2,500-1:10,000) using SuperSignal West Pico Chemiluminescent Substrate (Thermo Scientific) and developed using Blue Devil Autoradiography film (Genesee Scientific) or Bio-Rad Chemidoc XRS+ Imaging device.

### Extraction of metabolites for GC-MS analysis

For cell culture, polar metabolites and fatty acids were extracted using methanol/water/chloroform with [^2^H_31_]palmitate and norvaline as lipid and polar internal standards, respectively, and analyzed as previously described (22). Briefly, cells were washed twice with saline, quenched with -80C methanol and 4C water containing norvaline, scraped into Eppendorfs, and extracted with chloroform containing [^2^H_31_]Palmitate. After centrifugation, phases are dried separately. Samples are stored at -20C before analysis by GC-MS.

### GC-MS analysis

Dried polar metabolites were derivatized in 2% (w/v) methoxyamine hydrochloride (Thermo Scientific) in pyridine and incubated at 37°C for 60 min. Samples were then silylated with N-tertbutyldimethylsilyl-N-methyltrifluoroacetamide (MTBSTFA) with 1% tert-butyldimethylchlorosilane (tBDMCS) (Regis Technologies) at 45°C for 30 min. Polar derivatives were analyzed by GC-MS using a DB-35MS column (30m x 0.25mm i.d. x 0.25μm, Agilent J&W Scientific) installed in an Agilent 7890B gas chromatograph (GC) interfaced with an Agilent 5977A mass spectrometer (MS) with an XTR ion source. The dried lower chloroform phase was derivatized to form fatty acid methyl esters (FAMEs) via addition of 500μL 2% H_2_SO_4_ in MeOH and incubation at 50°C for 2h. FAMEs were extracted via addition of 100μL saturated salt solution and 2 500μL hexane washes. These were analyzed using a Select FAME column (100m × 0.25mm i.d.) installed in an Agilent 7890A GC interfaced with an Agilent 5975C MS using the following temperature program: 80°C initial, increase by 20°C/min to 170°C, increase by 1°C/min to 204°C, then 20°C/min to 250°C and hold for 10 min. The percent isotopologue distribution of each fatty acid and polar metabolite was determined and corrected for natural abundance using in-house algorithms adapted from Fernandez et al. (23). Mole percent enrichment (MPE) was calculated via the following equation:

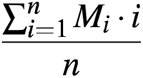

### Isotopomer spectral analysis (ISA) of fatty acids

Mass isotopomer distributions were determined by integrating metabolite ion fragments and correcting for natural abundance using in-house algorithms as previously described (14). The ISA compares a measured fatty acid isotopomer distribution to one that is simulated using a reaction network for palmitate synthesis, whereby 8 AcCoA molecules are consumed to form one palmitate molecule. Models were also generated for OCFA synthesis, whereby 1 PropCoA and 6-7 AcCoA molecules are consumed to form one OCFA as previously described (7). Parameters for the relative enrichment of the lipogenic AcCoA pool from a given [^13^C] tracer and the percentage of fatty acids that are *de novo* synthesized are extracted from a best-fit model using INCA v1.6 metabolic flux analysis software package (24). The 95% confidence intervals for both parameters were estimated by evaluating the sensitivity of the sum of squared residuals between measured and simulated fatty acid mass isotopomer distributions to small flux variations.

### Metabolic Flux Analysis

^13^C MFA was conducted using INCA, a software package based on elementary metabolite unit (EMU) framework (24, 25). Fluxes through the metabolic network comprising of glycolysis, the pentose phosphate pathway, the TCA cycle, BCAA catabolism, and fatty acid synthesis and oxidation were estimated by minimizing the sum of squared residuals between experimental MIDs and extracellular fluxes using nonlinear least squares regression for differentiated 3T3-L1 adipocytes traced with [U-^13^C_6_]glucose, [U-^13^C_5_]valine, [U-^13^C_6_]leucine. The best global fit was found after estimating 20 times using random initial guesses for all fluxes in the metabolic network. A χ^2^ statistical test was applied to assess the goodness-of-fit using α of 0.01. The 95% confidence intervals for each flux in the metabolic network were estimated by sensitivity analysis due to minor flux variations (26). See Supplementary Table 1 for metabolites used in the analysis, below for assumptions, and Supplementary Tables 2-3 for raw flux values.

^13^C metabolic flux analysis was conducted under the following assumptions:

- Cells were assumed to be at metabolic and isotopic steady state, i.e. intracellular free metabolite pool size and ^13^C enrichments are constant.
- Per well extracellular flux of glucose, lactate, glutamine, glutamate, alanine, valine, and leucine were assumed to be constant over the course of the labeling experiments.
- Experimental replicates from all three tracers were entered into INCA v2.0 and used simultaneously to estimate fluxes for sgControl and sgBckdha, resulting in tighter confidence intervals.
- Cells were assumed to be terminally differentiated and did not divert resources towards biomass synthesis.
- Succinate and fumarate are structurally symmetric and can be enzymatically converted to their subsequent products in either configuration.
- Labeled CO_2_ release during decarboxylation reactions is diluted upon release and does not reincorporate during carboxylation reactions.
- Separate mitochondrial and cytosolic pools of aspartate, oxaloacetate, malate, fumarate, pyruvate, acetyl-CoA, glutamate, and glutamine were modeled with exchange fluxes for malate, aspartate, glutamate, and glutamine.
- Relative branching of glucose flux to the oxidative pentose phosphate pathway (PPP) relative to the glucose uptake flux was set to 0.3%. (27)

### RNA isolation and quantitative RT-PCR analysis

Total RNA was purified from cultured cells using Trizol Reagent (Life Technologies) per the manufacturer’s instructions. First-strand cDNA was synthesized from 0.5μg of total RNA using High-capacity cDNA Reverse Transcription Kit with RNase inhibitor (Applied Biosystems) according to the manufacturer’s instructions. Individual 10μl SYBR Green real-time PCR reactions consisted of 2μL of diluted cDNA, 5μL of SYBR Green Supermix (Bio-Rad), and 1μL of primer master mix containing each forward and reverse primer at 5μM. For standardization of quantification, ribosomal RNA RPL27 was amplified simultaneously. The PCR was carried out on 96-well plates on a CFX Connect Real time System (Bio-Rad) using a three-stage program provided by the manufacturer: 95°C hot-start for 3 min, 40 cycles of 95°C for 10s and 60°C for 30s, followed by a 0.5°C/5 sec meltcurve generation protocol. Gene-specific primers used are listed in Supplementary Table 5.

### RNA-seq library preparation

Cells were lysed in Trizol and total RNA was extracted per manufacturer’s instructions. Stranded RNA-Seq libraries were prepared from polyA-enriched mRNA using the Illumina Stranded mRNA library prep kit. Library construction and sequencing was performed by the University of California San Diego (UCSD) Institute for Genomic Medicine. Libraries were run as PE100 on a NovaSeq S4 to a depth of ∼25 million reads.

### RNASeq Analysis and Gene Set Enrichment Analysis

The Kallisto/Sleuth differential expression pipeline analysis was performed for paired triplicate samples of NT, Bckdha, and Acad8 KO. The index was obtained from Pachter lab Kallisto website “Ensembl Transcriptomes v96”, mus_musculus.tar.gz. Kallisto was run for paired-end read quantification with sequence based bias correction (--bias) and 100 bootstraps (28). Normalized transcript abundances were further passed into sleuth using R, which were then aggregated to gene level (29). Ultimately, a pairwise Wald test was performed to compare sgControl v. sgBckdha and sgControl v. sgAcad8 using sleuth.

GSEA analysis using the KEGG pathway functional database was performed using WEB-based GEne SeT AnaLysis Toolkit (WebGestalt). A rank score was calculated through multiplication of the q-value with the sign of the fold change of sgBckdha and sgControl adipocytes. Only significantly differentially expressed genes (q<0.05) were included in the analysis. Redundancy reduction of the enriched gene sets was set to “All.” The MSigDB Hallmarks pathway functional database was downloaded in gene symbol format (http://www.gsea-msigdb.org/gsea/msigdb/collections.jsp) (30, 31). As this is a human pathway database, the differentially expressed genes in the present study were first mapped to their human homologs using the publicly available mouse human genome homology database compiled by the mouse genome informatics organization (http://www.informatics.jax.org/downloads/reports/) (32). GSEA analysis was then conducted using the WebGestalt platform, using the hallmarks pathway set as a custom database, and the human homologs of the differentially expressed genes (q<0.05) in this study and their associated rank score as the input gene list.

### Respirometry

Respirometry was conducted using a Seahorse XF96E Analyzer. For respiration studies, cells were plated at 5,000 cells/well and maintained in 2μg/mL puromycin-containing media until confluence was reached and differentiation was performed as described above. 8 days after differentiation was initiated, growth medium was replaced with DMEM (Sigma #5030) supplemented with 25mM HEPES, 8mM glucose, 2mM glutamine, and 1mM pyruvate. Mitochondrial respiration is calculated as the oxygen consumption rate sensitive to 1μM rotenone and 2μM antimycin A. Basal, maximal and ATP-linked respiration were calculated as previously described (33). Immediately after the experiment, cells were gently washed with PBS and either a) fixed with 4% paraformaldehyde and stained with CyQuant (Life Technologies) to determine cell number for normalization purposes or b) solubilized in RIPA buffer and quantified using BCA assay. Data from n=3 independent experiments is depicted after normalization within each experiment.

### Statistical analyses

Statistical analysis of RNA seq data was carried out as detailed above. Metabolomic data was analysed using MetaboAnalyst 5.0 where any missing data was input using k nearest neighbour, data was log-scaled and analysed using one-way ANOVA with post-hoc analysis using Tukey’s HSD. Significance was considered with a FDR <0.05. Data was autoscaled for visualization in heatmaps. All experiments were repeated at least 3 times and data from 1 representative experiment is shown unless other-wise specified. For analyses involving 2 groups, a two-tailed Student’s t-test was performed. For analyses with 3 groups or more, a one ANOVA was performed as appropriate with Dunnett’s multiple comparisons test using GraphPad Prism.

## Results

### *Bckdha* deficient adipocytes maintain lipid droplet formation but display a metabolically compromised transcriptional profile

To explore the impact of compromised BCAA catabolism in adipocytes, we generated polyclonal cultures of 3T3-L1 pre-adipocytes lacking *Bckdha* using CRISPR/Cas9 and then differentiated these using 3-isobutyl-1-methylxanthine (IBMX), dexamethasone, and insulin. BCKDHA was undetectable in these polyclonal cultures and we observed a 90%+ decrease in incorporation of [U-^13^C_6_]leucine in citrate (Fig. 1A-B), along with a decrease in BCAA uptake (Supp. Fig 1A). There was no apparent morphological difference with *Bckdha* deficiency, lipid droplet accumulation appeared unaffected (Fig 1C) expression of key adipocyte mRNA differentiation markers was largely unchanged in these polyclonal cultures (Supp. Fig. 1B). To further examine the impact of *Bckdha* deficiency, we performed RNAseq on our polyclonal cell populations. We found 887 genes that were significantly differentially expressed (Fig. 1D). The most significantly different genes included *F13a1* and *Sema3g*, which have previously been associated with adipogenesis and obesity (34, 35). To categorize these differentially expressed genes further, we used WEB-based gene set analysis toolkit (WebGestalt) to perform gene set enrichment analysis (GSEA) with the KEGG pathway functional database. We found seven significantly altered pathways with FDR <0.05, which comprised key functional, metabolic, and signaling pathways relevant for adipocyte function, including oxidative phosphorylation, thermogenesis, glycolysis, HIF-1 signaling, and AMPK signaling (Fig. 1E). We also performed GSEA using the MSigDB biological processes hallmark gene sets database. Notably, adipogenesis was the most downregulated pathway in this analysis, and epithelial-to-mesenchymal transition (EMT) was significantly upregulated (Fig. 1E), highlighting how *Bckdha* deficiency impacts cellular differentiation state even though lipid droplet accumulation is unchanged.

**Figure 1.**
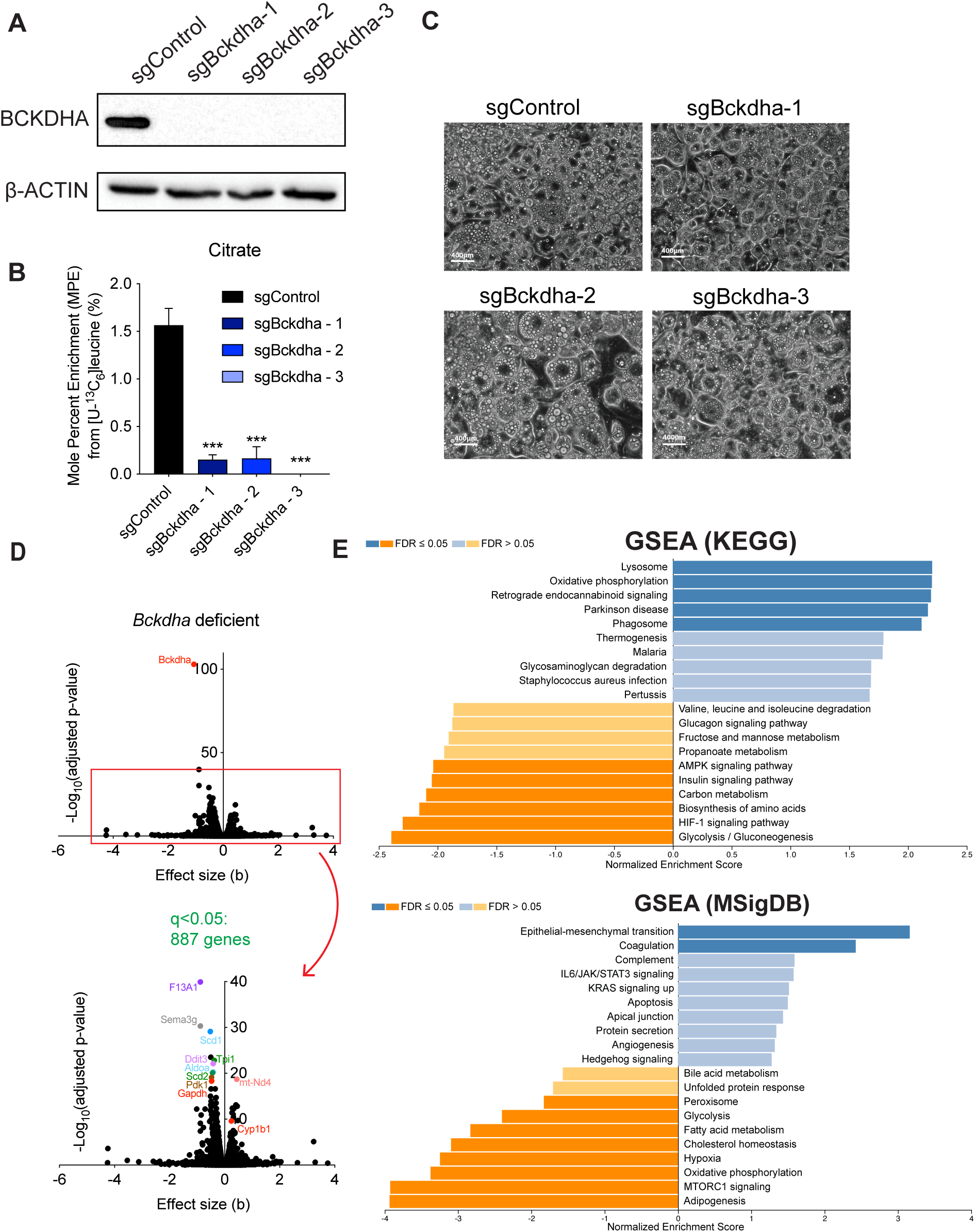
*Bckdha* deficient adipocytes maintain lipid droplet formation but display a metabolically compromised transcriptional profile. Characterization of sgControl and sgBckdha adipocytes 7 days post-induction of differentiation, A. Western blot of BCKDHA levels. B. brightfield images (scale bar = 400μm). C. Mole percent enrichment of citrate from [U-^13^C_6_]leucine (means +/- SEM, n=3, analysed via one-way ANOVA with Dunnetts multiple comparisons test, ***p<0.001.). D. Volcano plots of differentially expressed genes in *Bckdha*-deficient adipocytes compared to Control adipocytes. E. Gene set enrichment analysis of *Bckdha*-deficient adipocytes generated using WEB-based GEne SeT AnaLysis Toolkit (WebGestalt) (A-C). Results are depicted from one representative experiment which was repeated independently at least three times.

In addition to *Bckdha* deficiency, which prevents catabolism of all three BCAAs and downstream pathways, we also investigated the phenotype of adipocytes deficient in *Acad8*, which catalyzes the oxidation of valine-derived isobutyryl-CoA (and thus only blocks valine catabolism). ACAD8 protein knockdown (Supp. Fig. 1C) did not grossly affect lipid accumulation or adipogenic differentiation (Supp. Fig. 1D). In addition, RNAseq analysis of these *Acad8*-deficient adipocytes found only 65 differentially expressed genes (q<0.05) (Supp. Fig. 1E). Overall, these results suggest that *Acad8* deficiency and thus valine specific metabolism elicits more moderate effects compared to *Bckdha* deficiency and what has previously been reported for genes associated with leucine catabolism (11). Therefore, next we focused on characterizing how metabolism is altered in *Bckdha*-deficient adipocytes.

### *Bckdha*-deficient adipocytes reprogram central carbon metabolism

Glycolysis/gluconeogenesis was one of the most downregulated pathways in our GSEA of *Bckdha*-deficient adipocytes. To explore these changes more directly, we carried out intracellular metabolite analysis and quantified glucose uptake in Control and *Bckdha*-deficient adipocytes. Glucose uptake and lactate secretion were significantly decreased in *Bckdha*-deficient adipocytes (Fig. 2A-B) supporting the reduction in glycolysis suggested by our RNAseq analysis. In our intracellular metabolite analysis, pyruvate and TCA cycle intermediates along with the non-essential amino acids (NEAA) alanine, aspartate and glutamine were significantly decreased in *Bckdha*-deficient adipocytes (Fig. 2C and Supp. Fig. 2A). The primary metabolites increased were BCAAs, glutamate and BCKAs. To determine how impaired BCAA catabolism impacts the use of other carbon sources, we traced adipocytes with [U-^13^C_6_]glucose and [U-^13^C_5_]glutamine and quantified isotope enrichment in TCA intermediates. Importantly, we observed a significant increase in mole percent enrichment (MPE) of citrate, α-ketoglutarate (aKG), and succinate from [U-^13^C_6_]glucose in *Bckdha*-deficient adipocytes (Fig. 2D), with a relative increase in the M5 and M6 fractional amount in the isotopologue distribution of citrate which may indicate increased TCA cycle turnover (Supp. Fig. 2B). MPE from [U-^13^C_5_]glutamine was significantly higher in all TCA intermediates except succinate in *Bckdha*-deficient adipocytes (Fig. 2E). Analysis of oxygen consumption rates (OCR) in these cells in response to oligomycin indicated that the *Bckdha*-deficient cells have a significant decrease in ATP-linked respiration (Fig. 2F-G); however the basal and maximal OCR were not significantly affected (Supp. Fig. 2C-D). Collectively this indicates that ablation of BCAA catabolism in adipocytes is marked by compensatory increases in pyruvate and glutamine entry into the TCA cycle which minimizes the impact on mitochondrial respiration.

**Figure 2.**
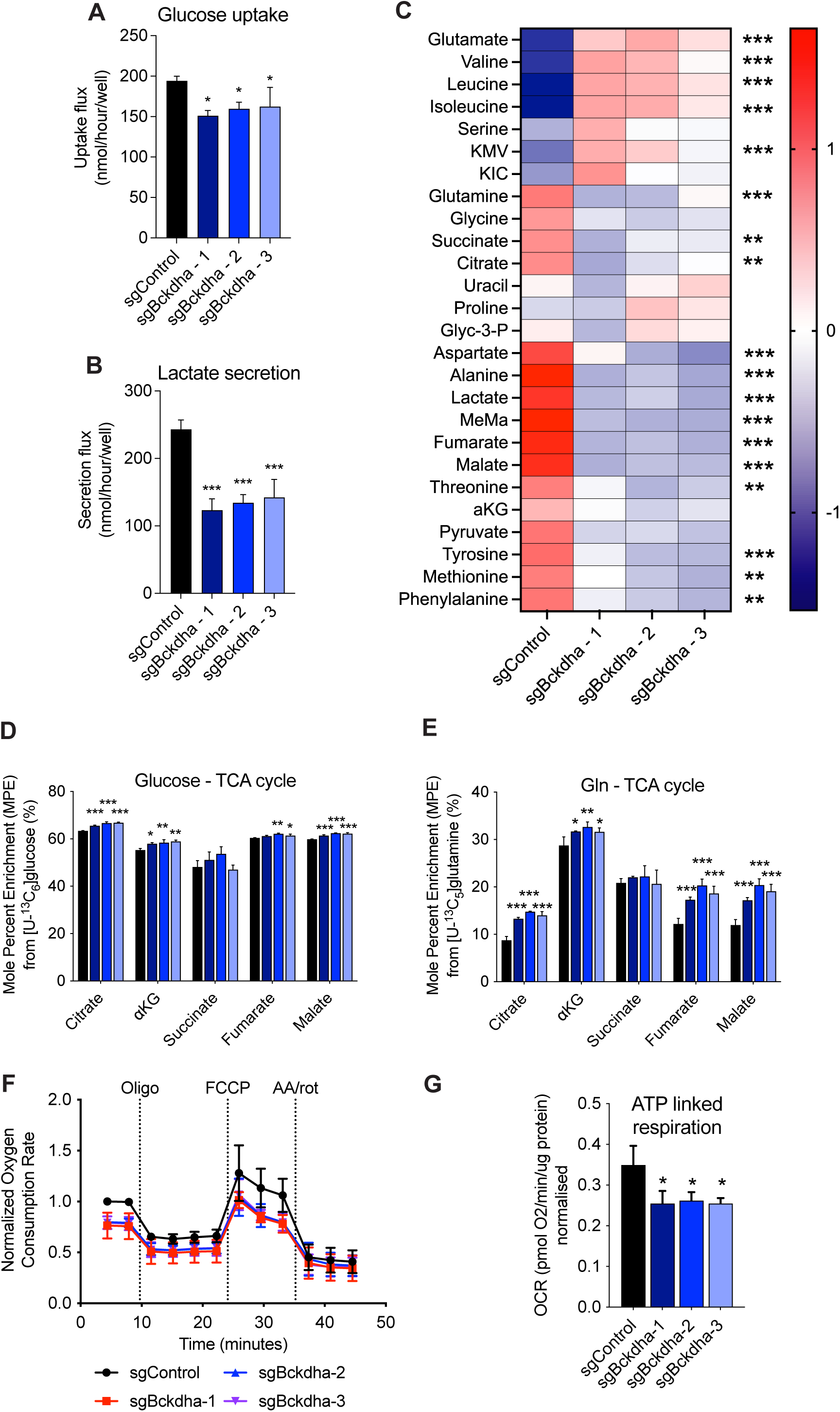
*Bckdha* deficient adipocytes reprogram central carbon metabolism. A. Molar amount of glucose uptake in Control or *Bckdha*-deficient adipocytes over 48 hours. B. Molar amount of lactate secretion in Control or *Bckdha*-deficient adipocytes over 48 hours. C. Heat map of the abundance of intracellular metabolites in *Bckdha*-deficient adipocytes, statistical analysis via one-way ANOVA (9 cellular replicates). FDR*<0.05, **<0.01, ***<0.001. D. Mole percent enrichment (MPE) of TCA cycle intermediates after 48 hours of incubation with [U-^13^C_6_]glucose. E. MPE of TCA cycle intermediates after 48 hours of incubation with [U-^13^C_5_]glutamine. F. Normalized oxygen consumption rate of 3T3-L1 Control or *Bckdha*-deficient adipocytes treated with the indicated pharmacological inhibitors (n=3 experiments internally normalized to sgControl). G. ATP linked respiration (n=3 individual experiments internally normalized to sgControl). (A-B, D-E) Data are presented as means ± SD with three cellular replicates. Results are depicted from one representative experiment which was repeated independently at least three times. (A-B, D-G) One-way ANOVA with Dunnett’s multiple comparisons test for each parameter. Significance in all is compared to sgControl. *p<0.05, **p<0.01, ***p<0.001.

Our metabolic studies were carried out under nutrient replete conditions; however, glucose utilisation in adipocytes is often functionally assessed via acute insulin-stimulated conditions in media depleted of nutrients other than glucose. To understand how glucose metabolism is changed in these conditions, we traced cells for 10 minutes in [U-^13^C_6_]glucose ± insulin following incubation in media depleted of serum and glucose (Fig. 3A). We found a general decrease in total ^13^C-labelled pyruvate and TCA intermediates under both basal(-insulin) and insulin stimulated conditions indicating a general decrease in glucose utilization is retained and potentially exasperated under these conditions (Fig. 3B-D). However, the fold change between basal and insulin stimulated was not decreased in *Bckdha*-deficient adipocytes (Fig. 3E) indicating altered glucose metabolism is unlikely to arise from a change in insulin signaling in these cells but rather a general decrease in glycolytic capacity.

**Figure 3.**
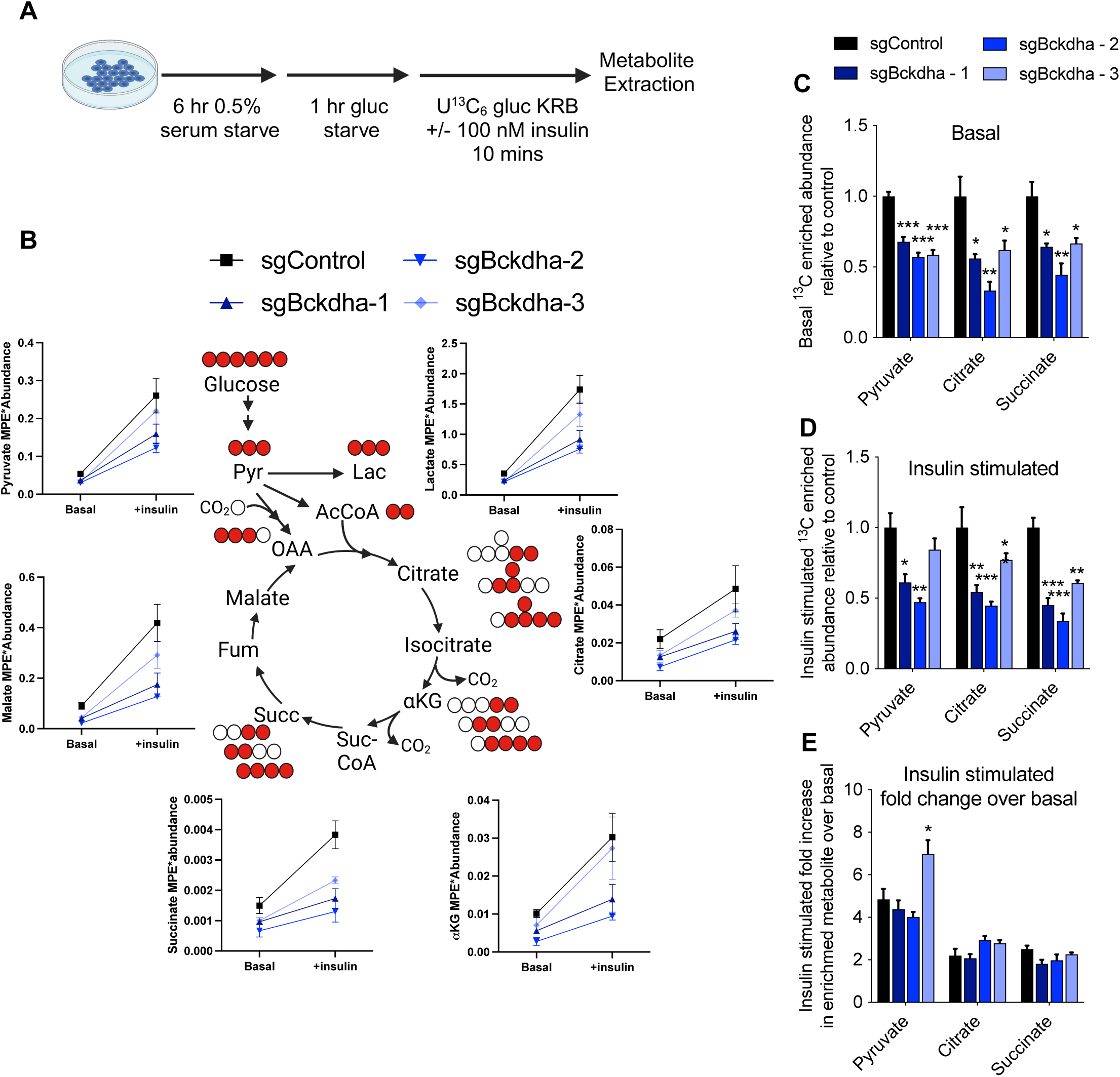
Insulin stimulated glucose uptake in *Bckdha* deficient cells. A. Schematic of experimental design. B-D. Levels of ^13^C enriched metabolite from [U-^13^C_6_]glucose in cells under basal and insulin stimulated conditions. E. Fold increase in ^13^C enriched metabolite bteween basal and insulin stimulated conditions. Data are presented as means ± SD with three cellular replicates. One-way ANOVA with Dunnett’s multiple comparisons test for each parameter. Significance in all is compared to sgControl. *p<0.05, **p<0.01, ***p<0.001.

### *Bckdha* deficiency alters lipogenic acetyl-CoA sourcing and fatty acid (FA) diversity

Given that lipid droplet accumulation was relatively unaffected by *Bckdha* deficiency and that our previous work demonstrated that BCAAs are a significant source of lipogenic acetyl-CoA in adipocytes (14), we sought to determine whether glucose utilization for *de novo* fatty acid synthesis was altered. Isotopomer spectral analysis (ISA) of palmitate labeling from [U-^13^C_6_]glucose demonstrated that the contribution of glucose derived carbon to the lipogenic acetyl-CoA pool was significantly increased despite the observed decrease in glucose uptake (Fig. 4A). 3T3-L1 adipocytes typically use BCAA-derived propionyl-CoA and branched-CoAs to synthesize odd chain fatty acids (OCFAs) and monomethyl branched-chain fatty acids (mmBCFAs) via FASN in addition to palmitate (7, 9, 14). As expected, OCFAs and mmBCFAs were greatly decreased, while the proportion of even-chain fatty acids was significantly increased in *Bckdha*-deficient adipocytes (Fig. 4B-C). ISA modeling also revealed a decrease in newly synthesized OCFAs and a modest increase in the proportion of newly synthesized C16:0 (Fig. 4D and Supp. Fig. 3A). However, the total amount of newly synthesized fatty acids (accounting for both percentage of pool newly synthesized and molar amount of FA) remained unchanged in *Bckdha*-deficient adipocytes (Fig. 4D). These results indicate that central carbon metabolism is rewired to sustain FA synthesis (i.e., acetyl-CoA sourcing) and overall abundances.

**Figure 4.**
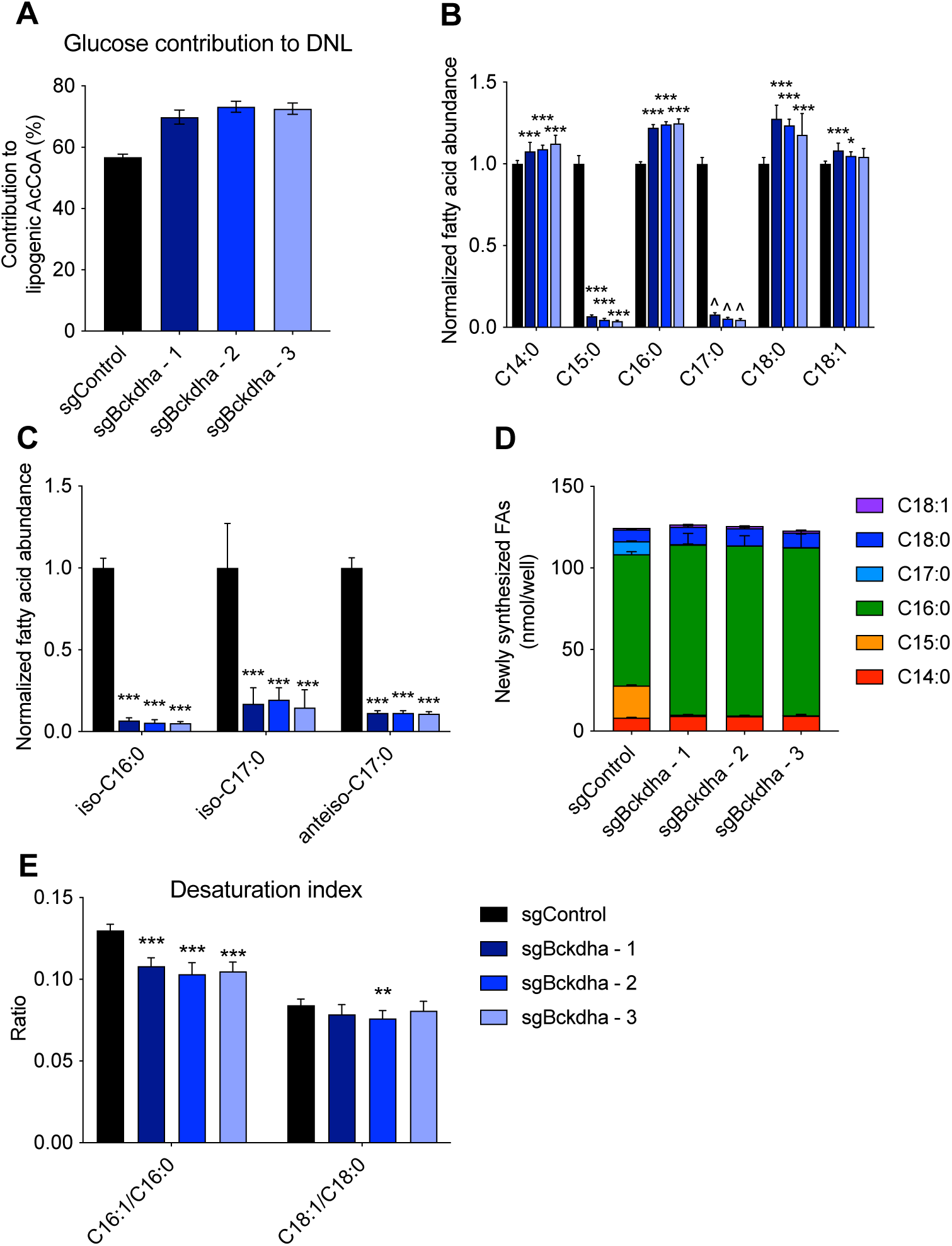
Bckdha deficiency alters lipogenic acetyl-CoA sourcing and fatty acid diversity. A. Contribution of [U-^13^C_6_]glucose to the lipogenic AcCoA pool used for *de novo* lipogenesis (DNL) as determined via ISA. B. Relative fold change in total odd-and even-straight-chain fatty acids in *Bckdha*-deficient 3T3-L1 adipocytes normalized to Control adipocytes. C. Relative fold change in total branched-chain fatty acids (BCFAs) in *Bckdha*-deficient 3T3-L1 adipocytes normalized to Control adipocytes. D. Molar amount of indicated fatty acids synthesized over 48 hours obtained by combining individual FA DNL values with pool size. E. Desaturation index in Control and *Bckdha*-deficient 3T3-L1 adipocytes. Data are presented as means ± SD (B-D), means with 95% confidence interval (C.I.) (A), and means ± SEM (E) with three cellular replicates. Each parameter was analysed via one-way ANOVA with Dunnett’s multiple comparisons testing except A, where significance is denoted as non-overlapping 95% C.I. Significance in all is compared to sgControl. Results are depicted from one representative experiment which was repeated independently at least three times. *p<0.05, **p<0.01, ***p<0.001.

Our RNAseq results also highlighted a decrease in the expression of the desaturase enzymes SCD1 and SCD2 in *Bckdha*-deficient adipocytes. To this end we calculated the desaturation index (ratio of monounsaturated to saturated FA) in each cell type. While the C18:1/C18:0 ratio was unchanged, the C16:1/C16:0 ratio was significantly decreased in *Bckdha-*deficient adipocytes indicating a decrease in C16:0 desaturation. As both C18:1 and C18:0 can be derived more from uptake than synthesis (compared to C16:1 and C16:0), this difference likely reflects a decrease in desaturation activity that is buffered by import of C18:1 from serum in media (Fig. 4E).

### MFA modeling of *Bckdha*-deficient adipocytes

As glucose uptake was decreased, yet utilization of glucose for *de novo* made fatty acids was increased, we developed a ^13^C metabolic flux analysis (^13^C-MFA) model incorporating central carbon metabolism and lipid biosynthesis to better understand how glucose metabolism was being altered. We incorporated data from Control and *Bckdha*-deficient adipocytes cultured with either [U-^13^C_6_]glucose, [U-^13^C_6_]leucine, or [U-^13^C_5_]valine for 48 hours into a MFA model using INCA (24). The network encompassed relevant reactions within glycolysis, TCA metabolism, BCAA oxidation, and fatty acid synthesis, with compartmentalization of several pathways included (Fig. 5). Isotope enrichment data for pyruvate, TCA cycle intermediates, intracellular and extracellular glutamine, intracellular glutamate, leucine, KIC, valine, and fatty acids C15:0, C16:0, and C17:0, as appropriate were included in the model (Supplementary Table 1) along with direct measurements of uptake and secretion fluxes for glucose, lactate, alanine, valine, leucine, glutamate, and glutamine (Supplementary Table 2-3).

**Figure 5.**
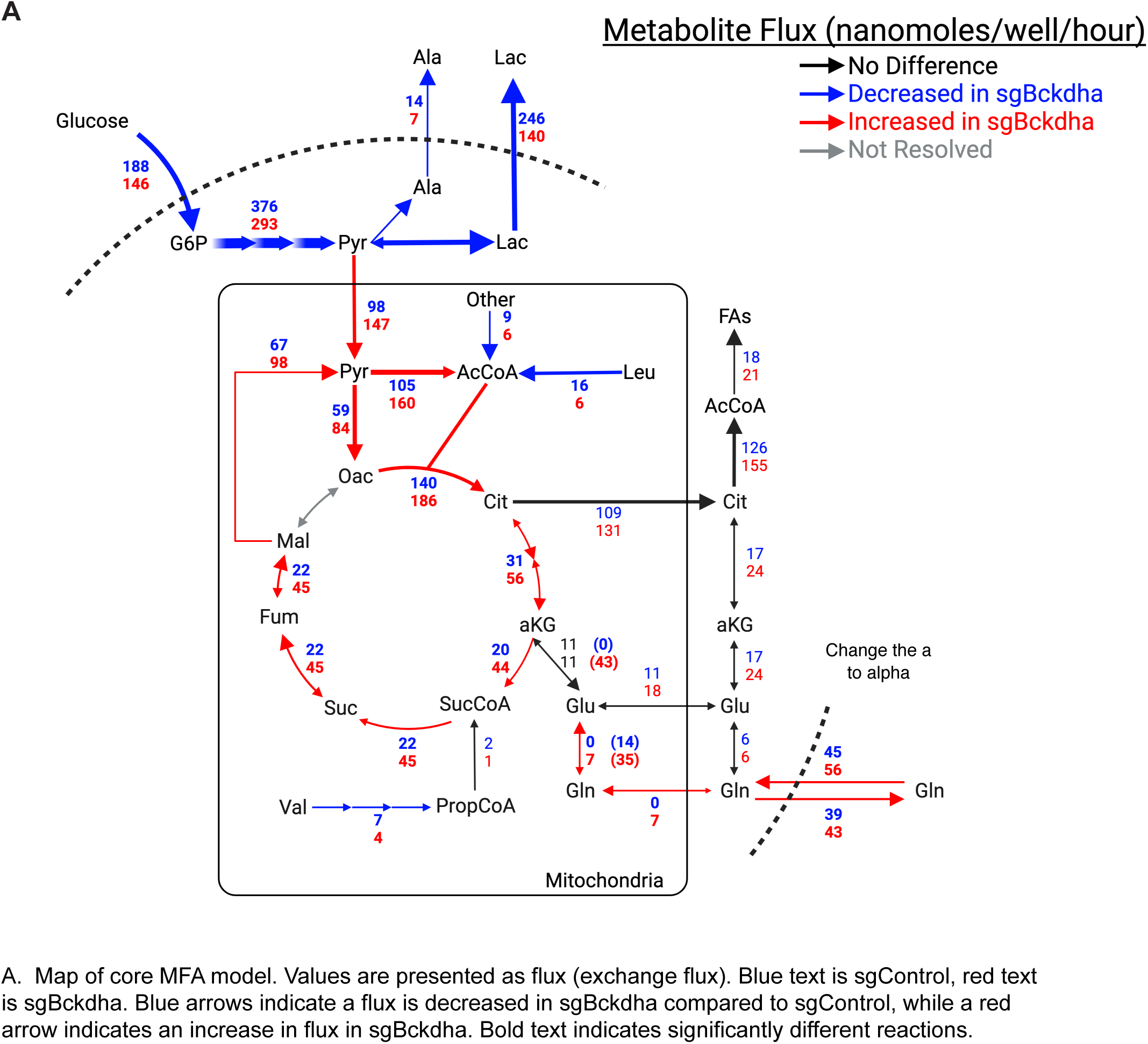
Metabolic flux analysis reveals significant reprogramming of glycolysis and TCA metabolism. Map of core MFA model. Values are presented as net flux (exchange flux). Blue text is sgControl, red text is sgBckdha. Blue arrows indicate a flux is decreased in sgBckdha compared to sgControl, while a red arrow indicates an increase in flux in sgBckdha. Bold text indicates significantly different reactions.

Flux estimates and confidence from the model highlighted a significant decrease in glucose uptake and both lactate and alanine secretion in 3T3-L1 *Bckdha*-deficient adipocytes. On the other hand, *Bckdha*-deficient adipocytes exhibited a marked increase in pyruvate dehydrogenase (PDH) and pyruvate carboxylase (PC) flux which is consistent with the observed increase in glucose enrichment in TCA intermediates and glucose derived carbon being used for fatty acid synthesis when acetyl-CoA derived from leucine/isoleucine catabolism is decreased. Indeed, the PDH kinase PDK1 was one of the most significantly downregulated genes in our RNAseq analysis (Fig. 1D), indicating there may be decreased inhibitory regulation of PDH in *Bckdha*-deficient adipocytes. We also observed increased pyruvate cycling and although we could not resolve cytosolic from mitochondrial malic enzyme flux, these results highlight the importance of pyruvate cycling in these cells, as cytosolic malic enzyme is a major producer of lipogenic NADPH in adipocytes (27, 36).

Notably, anaplerotic flux of BCAAs to succinyl-CoA was low in all cells, presumably due to the lack of vitamin B12 present in these cultures (14). This shift in carbon balancing from BCAA oxidation to glucose/pyruvate oxidation resulted in the model predicting an increase in oxidative TCA cycle flux. Indeed, glutamine contribution to the TCA cycle was also estimated to be increased in *Bckdha*-deficient adipocytes, which also exhibited dramatically higher rates of exchange between glutamate and a-KG. These results are in agreement with our independent results using [U-^13^C_5_]glutamine as a tracer (Fig. 2E), which were not incorporated into the model due to the high exchange with medium glutamine. Notably, the model predicted both an increase in the uptake (56 nmol/well/h vs 45 nmol/well/h) and secretion (43 nmol/well/h vs 39 nmol/well/h) of glutamine resulting in a net decrease in total glutamine in spent medium. Overall, these findings highlight the broad impact of *Bckdha* deficiency on both central carbon and nitrogen metabolism (37). In this manner, altered BCAA metabolism can have broad impacts on adipose tissue function associated with carbohydrate disposal and amino acid homeostasis.

## Discussion

Here we explored how the absence of BCAA or valine catabolism impacts differentiation and metabolism in cultured 3T3-L1 adipocytes. In contrast to previous studies, we found that adipocyte lipid droplet accumulation and *de novo* lipogenesis are maintained in the absence of BCKDH. However central carbon and non-essential amino acid synthesis are re-wired such that glycolysis and lactate secretion are potently downregulated and there is increased use of pyruvate and glutamine to fuel TCA cycle metabolism and *de novo* lipogenesis. Overall, our findings indicate that decreased BCAA catabolism during adipogenesis has subtle impacts on the differentiation state and metabolic capacity of adipocytes which may prime the cells for further dysfunction in the obese state and impact intercellular metabolite cross-talk in adipose tissue.

The most notable feature of metabolism in our *Bckdha*-deficient adipocytes was a decrease in glycolysis and lactate secretion under both typical nutrient-replete conditions when cells were monitored over 24 hours and acute insulin stimulation. However, the fold difference in glucose utilisation between basal and insulin-stimulated conditions was not decreased indicating insulin signalling is not compromised but rather there is a general decrease in glycolytic capacity. This metabolic change was also reflected in the gene expression signature where glycolysis was the most downregulated pathway in our GSEA using the KEGG functional database. This suggests there is cross-talk between BCAA catabolism and signaling mechanisms necessary to regulate glycolysis during adipocyte differentiation. GSEA also indicated a change in both mTORC1 and HIF-1 signaling which are known to regulate glycolysis (38). Although pathological HIF-1 accumulation has been shown to inhibit adipocyte differentiation (25), previous studies have also found that HIF-1 alpha can support glycolytic flux in brown adipocytes (39). The potential functional impact of this in an adipose tissue context is unclear, however recent studies have suggested lactate production is prioritized by adipocytes (40). In addition, lactate is emerging as an important substrate for fueling TCA cycle metabolism (41) and inter-cellular crosstalk (42). Overall, further studies will be required to understand cross-talk between BCAA catabolism and glycolysis during adipocyte differentiation and whether suppression of BCAA catabolism during adipogenesis *in vivo* would impact adipose lactate metabolism.

Surprisingly, although glucose uptake was decreased, our data indicate a significant increase in pyruvate utilisation via increased flux through both pyruvate dehydrogenase and pyruvate carboxylase. Previous studies have demonstrated the presence of significant crosstalk between pyruvate and BCAA or BCKA metabolism. Indeed, inhibition of the mitochondrial pyruvate carrier in human hepatoma cell lines results in decreased inhibitory phosphorylation of BCKDH indicating altered pyruvate utilisation can result in a compensatory increase in BCAA breakdown (43). Our data suggests that similar cross-talk occurs here where decreased BCAA catabolism results in a compensatory increase in pyruvate utilization. However, elevated BCKAs due to decreased BCKDH activity or via exogenous addition of BCKAs have previously been shown to inhibit pyruvate dehydrogenase in hepatocytes, heart and the brain (44–47) as well as the mitochondrial pyruvate carrier in hepatocytes (48). While our data indicates that the opposite occurs here where decreased BCKDH activity enhances pyruvate utilisation, our BCKA accumulation in this context was subtle and only significant for KMV. This may be due to a concurrent decrease in BCAT2 activity which has been noted to occur with altered BCKDH activity (49) resulting in greater BCAA accumulation rather than BCKAs which can protect against this phenomenon (48). In addition, BCKDH KO was accompanied by a decrease in BCAA uptake indicating compensatory mechanisms in cell culture. Thus, in contrast to other cell types, adipocyte pyruvate utilization is not compromised by decreased BCKDH activity but rather enhanced which allows the cell to maintain TCA flux and *de novo* lipogenesis.

*Bckdha-*deficient adipocytes also displayed a significant difference in their NEAA levels and fatty acid composition. Of note, net glutamine levels in the media significantly decreased due to increased utilization in the TCA cycle which was not compensated by *de novo* glutamine synthesis, which is generally elevated in adipocytes (14). Decreased glutamine levels in obese adipose tissue have previously been noted and associated with inflammation and adipose dysfunction (50). *Bckdha*-deficient adipocytes also had decreased mmBCFAs and OCFAs which we have previously shown are synthesized from intermediates in the BCAA catabolic pathway (7). mmBCFAs have been proposed to signal via mTORC1 (51) and have been implicated in altered immune cell signaling (52, 53) thus altered levels of these metabolites may have a significant impact on intercellular communication with other adipose resident cells *in vivo*.

Finally, we found that decreased BCAA catabolism had a subtle effect on overall markers of adipocyte differentiation. Lipid droplet accumulation was not impacted in our polyclonal *Bckdha*-deficient adipocytes and this contrasts our prior results targeting *Bckdha* using shRNA (14) and those of others utilizing shRNA or siRNA to alter MCCC1 and HIBCH levels (11, 19). However, in our RNAseq analysis using the molecular signature database, adipogenesis was one of the most downregulated pathways along with an increase in expression of genes associated with epithelial-to-mesenchymal transition indicating a more fibroblast-like gene signature. Notably, *Bckdha* deficiency induced much broader changes in metabolism and gene expression than *Acad8* deficiency, where valine-mediated anaplerosis was reduced without grossly altering intermediary metabolism or gene expression.

Overall, our studies highlight how *Bckdha* deficiency during differentiation re-wires the metabolic network and the levels of multiple metabolites that have been shown to play a role in intracellular cross-talk *in vivo*. However, further studies are needed at multiple time-points to understand how altered BCAA catabolism during adipocyte differentiation impacts the signaling networks that regulate transcription of other metabolic pathways and the key nodes and timepoints where it is most influential.

## Acknowledgments

We thank all members of the Metallo laboratory for support and helpful discussions. We would also like to thank Dr. Yan Wu for guidance in RNAseq analysis.

## Author Contributions

C.R.G., C.M.M. and M.W. conceived and designed the study, analysed and interpreted data and wrote the manuscript with help from all other authors. K.W.-R. performed MFA analysis. R.T. conducted MSigDB-based RNAseq analysis. J.D.H. assisted with gene expression analysis, stable-isotope tracing experiments and GC-MS analysis. A.N.M. provided Seahorse equipment and related training and advice. C.R.G. performed all other experiments.

## Supplementary Figures

**Supplementary figure 1.**
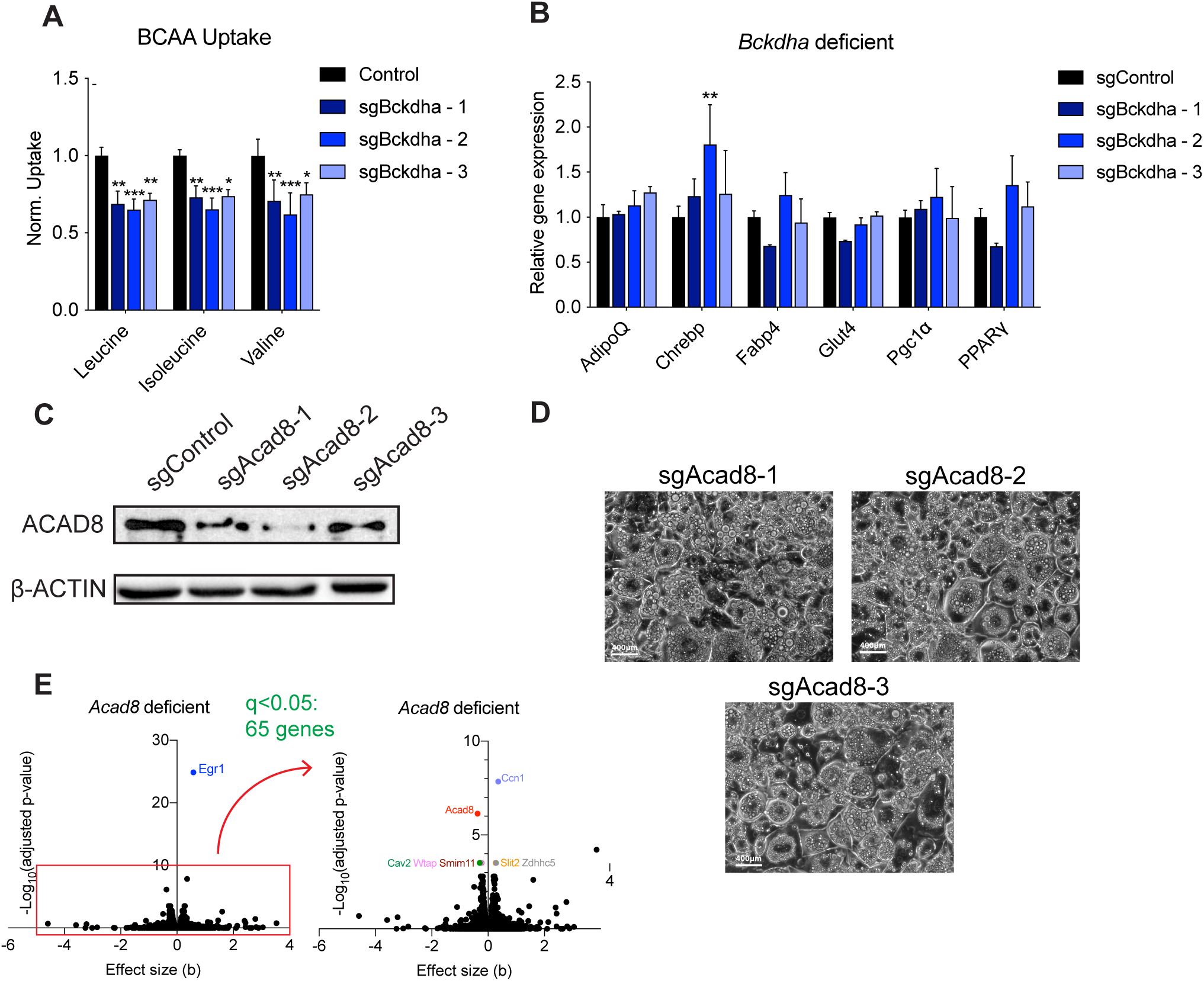
A. Normalized relative uptake of leucine, isoleucine, and valine in Control and *Bckdha*-deficient adipocytes. B. Expression of adipocyte differentiation markers normalized to sgControl. C. Western blot of ACAD8 and β-ACTIN. D. Brightfield images of differentiated Control and sgAcad8 adipocytes 7 days post-induction of differentiation (scale bar = 400μm). E. Volcano plots of differentially expressed genes in *Acad8*-deficient adipocytes compared to Control adipocytes. Data are presented as means ± SD (A-B) with three cellular replicates. Each parameter was analysed via one-way ANOVA with Dunnett’s multiple comparisons testing. Significance in all is compared to sgControl. Results are depicted from one representative experiment which was repeated independently at least three times. *p<0.05, **p<0.01, ***p<0.001.

**Supplementary Figure 2.**
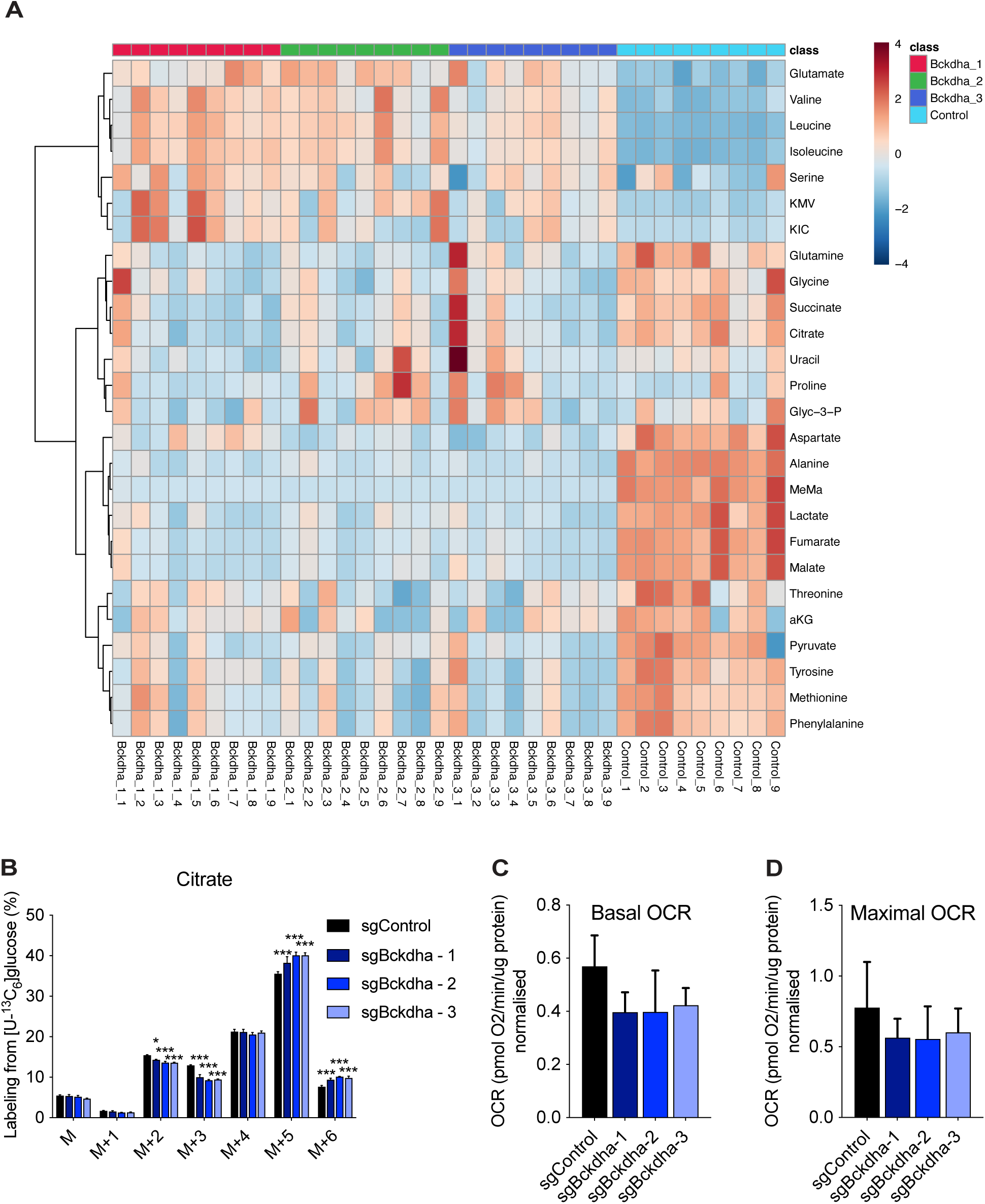
A. Heat map of the abundance of intracellular metabolites in *Bckdha*-deficient adipocytes showing each replicate sample. B. Citrate mass isotopomer distribution (MID) after 48 hours in [U-^13^C_6_]glucose. Results are depicted from one representative experiment which was repeated independently at least three times. Each parameter was analysed via one-way ANOVA with Dunnett’s multiple comparisons testing. C-D. Basal and maximal oxygen consumption rates in Bckdha deficient adipocytes. n=3 experiments internally normalized to sgControl, analysed via one-way ANOVA with Dunnett’s multiple comparisons testing. *p<0.05, **p<0.01, ***p<0.001.

**Supplementary figure 3.**
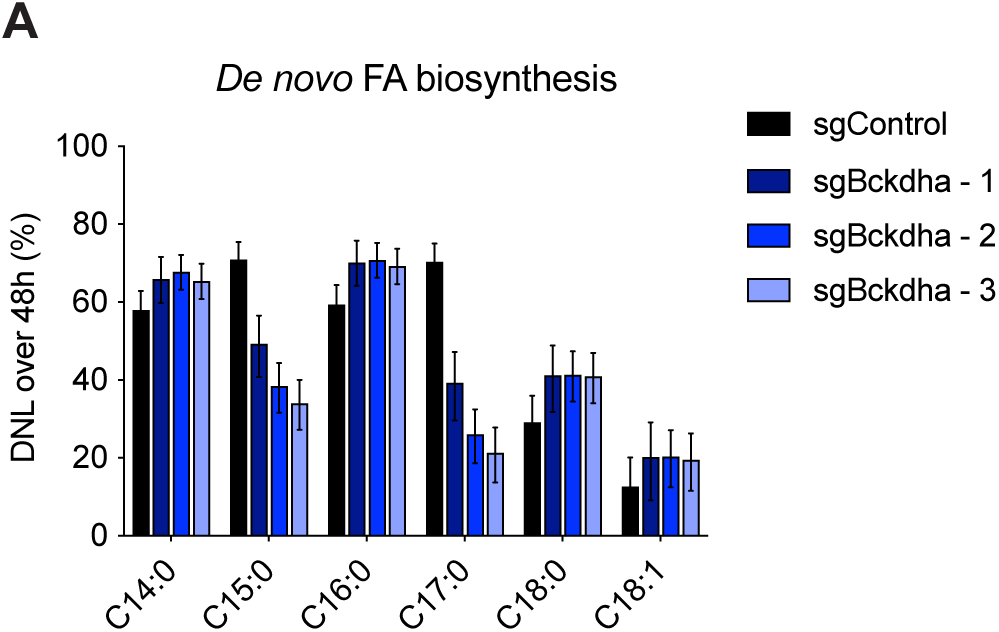
A. Percent of fatty acids derived from *de novo* lipogenesis in Control and *Bckdha*-deficient 3T3-L1 adipocytes over 48 hours obtained via isotopomer spectral analysis (ISA). Data is shown as means with 95% confidence interval (C.I.). Significance is denoted as non-overlapping 95% C.I.

