## Supplemental Tables for "Impaired BCAA catabolism during adipocyte differentiation decreases glycolytic flux"

**Supplementary Table 1**: GC-MS metabolite fragment ions. Metabolites used in sgControl v. sgBckdha MFA model:

Metabolite fragment ions used for GC-MS analysis:

| **Metabolite** | **Carbons** | **Formula** | **Mass (m/z)** |
|---|---|---|---|
| Pyruvate | 123 | $C_6H_{12}O_3NSi$ | 174 |
| aKG | 12345 | $C_{14}H_{28}O_5NSi_2$ | 346 |
| Fumarate | 1234 | $C_{12}H_{23}O_4Si_2$ | 287 |
| Malate | 1234 | $C_{18}H_{39}O_5Si_3$ | 419 |
| Glutamate | 12345 | $C_{19}H_{42}O_4NSi_3$ | 432 |
| Glutamine | 12345 | $C_{19}H_{43}O_3N_2Si_3$ | 431 |
| Citrate | 123456 | $C_{20}H_{39}O_6Si_3$ | 459 |
| Palmitate | 1-16 | $C_{17}H_{34}O_2$ | 270 |
| Pentadecanoic Acid | 1-15 | $C_{16}H_{32}O_2$ | 256 |
| Heptadecanoic Acid | 1-17 | $C_{18}H_{36}O_2$ | 284 |
| Leucine | 23456 | $C_{13}H_{32}O_1N_1Si_2$ | 274 |
| Ketoisocaproate | 123456 | $C_9H_{18}O_3N_1Si_1$ | 216 |
| Valine | 12345 | $C_{13}H_{30}O_2N_1Si_2$ | 288 |

**Supplementary Table 2**. MFA of 3T3-L1 sgControl. Full results of the $^{13}$C-MFA model.

| 3T3-L1 sgControl Pathway/Reaction | Flux (nanomoles/well/hr) | 95% Confidence Interval Lower Bound | Upper Bound |
|---|---|---|---|
| **Glycolysis** | | | |
| Glc.x -> G6P | 188.03 | 184.41 | 191.68 |
| G6P <-> F6P | 187.43 | 183.81 | 191.08 |
| G6P <-> F6P | 94.24 | 0.00 | Inf |
| F6P -> DHAP + GAP | 187.83 | 184.21 | 191.48 |
| DHAP <-> GAP | 187.83 | 184.21 | 191.48 |
| DHAP <-> GAP | 0.00 | 0.00 | Inf |
| GAP <-> 3PG | 375.86 | 368.62 | 383.17 |
| GAP <-> 3PG | 86.70 | 0.00 | Inf |
| 3PG -> PEP | 375.86 | 368.62 | 383.17 |
| PEP -> Pyr.c | 375.86 | 368.62 | 383.17 |
| Pyr.c <-> Lac | 264.19 | 254.49 | 273.68 |
| Pyr.c <-> Lac | 2.86 | 0.00 | Inf |
| Lac -> Lac.x | 264.19 | 254.49 | 273.68 |
| Pyr.c -> Ala | 13.97 | 8.98 | 15.58 |
| Pyr.m -> Ala | 0.00 | 0.00 | 4.88 |
| Ala -> Ala.x | 13.97 | 12.35 | 15.58 |
| **Pentose Phosphate Pathway** | | | |
| G6P -> P5P + CO2 | 0.60 | 0.57 | 0.60 |
| P5P + P5P <-> S7P + GAP | 0.20 | 0.19 | 0.20 |
| P5P + P5P <-> S7P + GAP | 0.00 | 0.00 | NaN |
| S7P + GAP <-> F6P + E4P | 0.20 | 0.19 | 0.20 |
| S7P + GAP <-> F6P + E4P | 0.00 | 0.00 | Inf |
| P5P + E4P <-> F6P + GAP | 0.20 | 0.19 | 0.20 |
| P5P + E4P <-> F6P + GAP | 0.00 | 0.00 | Inf |
| **Anaplerotic Reactions** | | | |
| Pyr.c -> Pyr.m | 97.71 | 86.04 | 109.92 |
| Pyr.m + CO2 -> Oac.m | 58.99 | 51.32 | 67.54 |
| Oac.c -> PEP + CO2 | 0.00 | 0.00 | 2.71 |
| Mal.m -> Pyr.m + CO2 | 66.51 | 58.13 | 75.78 |
| Mal.c -> Pyr.c + CO2 | 0.00 | 0.00 | 2.67 |
| PropCoA + CO2 -> SucCoA | 1.66 | 0.95 | 2.45 |
| Gln.c <-> Glu.c | 6.29 | 2.17 | 6.29 |
| Gln.c <-> Glu.c | 0.00 | 0.00 | 4.12 |
| Glu.c -> Glu.x | 0.27 | 0.20 | 0.34 |
| Glu.c <-> Glu.m | -11.06 | -16.15 | -8.09 |

| | | | |
|---|---|---|---|
| Glu.c <-> Glu.m | 0.00 | 0.00 | 4.12 |
| Glu.m <-> Akg.m | -11.22 | -12.85 | -8.32 |
| Glu.m <-> Akg.m | 0.00 | 0.00 | 5.15 |
| Akg.c <-> Glu.c | -17.08 | -19.00 | -14.11 |
| Akg.c <-> Glu.c | 1240.00 | 0.00 | Inf |
| Akg.m <-> Akg.c | 0.00 | -2.17 | 0.90 |
| Akg.m <-> Akg.c | 0.00 | 0.00 | 0.90 |
| Akg.c + CO2 <-> Cit.c | 17.08 | 14.12 | 18.90 |
| Akg.c + CO2 <-> Cit.c | 0.00 | 0.00 | 1.04 |
| Glu.m <-> Gln.m | 0.16 | -3.97 | 0.62 |
| Glu.m <-> Gln.m | 14.44 | 12.05 | 17.57 |
| Gln.x -> Gln.e | 6.13 | 5.67 | 6.59 |
| Gln.e -> Gln.c | 38.92 | 38.42 | 39.42 |
| Gln.c -> Gln.e | 32.79 | 32.52 | 33.05 |
| Gln.m <-> Gln.c | 0.16 | -3.97 | 0.62 |
| Gln.m <-> Gln.c | 54900.00 | 11.98 | Inf |
| **TCA Cycle** | | | |
| Pyr.m -> AcCoA.m + CO2 | 105.22 | 92.85 | 118.16 |
| AcCoAOther -> AcCoA.m | 18.48 | 14.99 | 22.35 |
| AcCoA.m + Oac.m -> Cit.m | 140.10 | 122.65 | 158.49 |
| Cit.m <-> Akg.m + CO2 | 31.20 | 27.33 | 35.10 |
| Cit.m <-> Akg.m + CO2 | 0.00 | 0.00 | 0.79 |
| Akg.m -> SucCoA + CO2 | 19.97 | 17.06 | 23.28 |
| SucCoA -> Suc | 21.63 | 18.25 | 25.45 |
| Suc <-> Fum.m | 21.63 | 18.25 | 25.45 |
| Suc <-> Fum.m | 0.00 | 0.00 | Inf |
| Fum.m <-> Mal.m | 21.63 | 18.25 | 25.45 |
| Fum.m <-> Mal.m | 553.87 | 0.00 | Inf |
| Mal.m <-> Oac.m | 653.51 | -1260.00 | 1440.00 |
| Mal.m <-> Oac.m | 0.00 | 0.00 | 2010.00 |
| Oac.m <-> Asp.m | 572.40 | -1440.00 | 1360.00 |
| Oac.m <-> Asp.m | 398.54 | 0.00 | Inf |
| Mal.c <-> Oac.c | 130.53 | -1400.00 | Inf |
| Mal.c <-> Oac.c | 0.00 | 0.00 | Inf |
| Oac.c <-> Asp.c | 256.52 | -1360.00 | Inf |
| Oac.c <-> Asp.c | 533.54 | 0.00 | Inf |
| Asp.c -> Fum.c | 828.92 | 0.00 | Inf |
| Mal.c <-> Fum.c | -828.92 | -Inf | 0.00 |
| Mal.c <-> Fum.c | 1580.00 | 0.00 | Inf |
| Mal.c <-> Mal.m | 698.39 | -1210.00 | 1390.00 |

| | | | |
|---|---|---|---|
| Mal.c <-> Mal.m | 86100.00 | 0.00 | Inf |
| Asp.m <-> Asp.c | 572.40 | -1440.00 | 1360.00 |
| Asp.m <-> Asp.c | 0.00 | 0.00 | Inf |
| **BCAA Catabolism** | | | |
| Leu.x -> Leu | 9.37 | 9.05 | 9.69 |
| Leu <-> KIC | 5.47 | 4.71 | 6.29 |
| Leu <-> KIC | 0.34 | 0.00 | Inf |
| KIC <-> IsoVCoA + CO2 | 5.47 | 4.71 | 6.29 |
| KIC <-> IsoVCoA + CO2 | 0.17 | 0.07 | 0.65 |
| IsoVCoA + CO2 <-> HMGCoA | 5.47 | 4.71 | 6.29 |
| IsoVCoA + CO2 <-> HMGCoA | 0.17 | 0.00 | 1500000.00 |
| HMGCoA -> AcCoA.m + AcCoA.m + AcCoA.m | 5.47 | 4.71 | 6.29 |
| LeuT -> Leu | 3.93 | 3.74 | 4.13 |
| Leu -> LeuP | 7.84 | 6.90 | 8.73 |
| Val.x -> Val | 5.85 | 4.96 | 6.74 |
| ValT -> Val | 1.51 | 1.26 | 1.77 |
| Val -> ValP | 0.00 | 0.00 | 2.57 |
| Val <-> IsoBCoA + CO2 | 7.36 | 5.16 | 8.48 |
| Val <-> IsoBCoA + CO2 | 0.54 | 0.38 | 0.72 |
| IsoBCoA <-> MMA | 7.35 | 4.89 | 8.48 |
| IsoBCoA <-> MMA | 0.00 | 0.00 | Inf |
| MMA -> PropCoA + CO2 | 7.35 | 4.89 | 8.48 |
| **Fatty Acid Synthesis** | | | |
| Cit.m -> Cit.c | 108.91 | 93.79 | 125.05 |
| Cit.c -> AcCoA.c + Oac.c | 125.99 | 110.17 | 142.64 |
| AcCoA.c + AcCoA.c + AcCoA.c + AcCoA.c + AcCoA.c + AcCoA.c + AcCoA.c + AcCoA.c -> Palm.s | 10.62 | 7.97 | 17.15 |
| Palm.s -> Palm | 10.62 | 7.97 | 17.15 |
| Palm.d -> Palm | 7.36 | 5.47 | 12.08 |
| PropCoA + AcCoA.c + AcCoA.c + AcCoA.c + AcCoA.c + AcCoA.c + AcCoA.c -> C15FA.s | 5.69 | 1.48 | 7.04 |
| C15FA.s -> C15FA | 5.69 | 1.48 | 7.04 |
| C15FA.d -> C15FA | 1.16 | 0.00 | 1.56 |
| C15FA + AcCoA.c -> C17FA.s | 6.85 | 1.01 | 8.49 |
| C17FA.s -> C17FA | 6.85 | 1.01 | 8.49 |
| C17FA.d -> C17FA | 2.22 | 0.00 | 2.82 |

| | | | |
|---|---|---|---|
| IsoBCoA + AcCoA.c + AcCoA.c + AcCoA.c + AcCoA.c + AcCoA.c + AcCoA.c + AcCoA.c -> IsoC16.s | 0.01 | 0.00 | 2.51 |
| IsoC16.s -> IsoC16 | 0.01 | 0.00 | 2.51 |
| IsoC16.d -> IsoC16 | 0.00 | 0.00 | 1.46 |
| **Dilution/Mixing** | | | |
| 0*Pyr.c -> Pyr.mnt | 0.00 | 0.00 | 0.01 |
| 0*Pyr.m -> Pyr.mnt | 1.00 | 0.99 | 1.00 |
| 0*Mal.c -> Mal.mnt | 0.00 | 0.00 | 1.00 |
| 0*Mal.m -> Mal.mnt | 1.00 | 0.00 | 1.00 |
| 0*Asp.c -> Asp.mnt | 0.98 | 0.00 | 1.00 |
| 0*Asp.m -> Asp.mnt | 0.02 | 0.00 | 1.00 |
| 0*Fum.m -> Fum.mnt | 0.99 | 0.00 | 1.00 |
| 0*Fum.c -> Fum.mnt | 0.01 | 0.00 | 1.00 |
| 0*Cit.m -> Cit.mnt | 0.76 | 0.53 | 0.97 |
| 0*Cit.c -> Cit.mnt | 0.24 | 0.03 | 0.47 |
| 0*Glu.m -> Glu.mnt | 1.00 | 0.74 | 1.00 |
| 0*Glu.c -> Glu.mnt | 0.00 | 0.00 | 0.26 |
| 0*Gln.m -> Gln.mnt | 0.00 | 0.00 | 1.00 |
| 0*Gln.c -> Gln.mnt | 1.00 | 0.00 | 1.00 |
| 0*Akg.m -> Akg.mnt | 0.46 | 0.41 | 0.55 |
| 0*Akg.c -> Akg.mnt | 0.54 | 0.45 | 0.59 |
| Pyr.mnt -> Pyr.fix | 1.00 | 1.00 | 1.00 |
| Asp.mnt -> Asp.fix | 1.00 | 1.00 | 1.00 |
| Mal.mnt -> Mal.fix | 1.00 | 1.00 | 1.00 |
| Fum.mnt -> Fum.fix | 1.00 | 1.00 | 1.00 |
| Cit.mnt -> Cit.fix | 1.00 | 1.00 | 1.00 |
| Akg.mnt -> Akg.fix | 1.00 | 1.00 | 1.00 |
| Glu.mnt -> Glu.fix | 1.00 | 1.00 | 1.00 |
| Gln.mnt -> Gln.fix | 1.00 | 1.00 | 1.00 |
| SSR | 774 | 637 | 785 |

**Supplementary Table 3**. MFA of 3T3-L1 sgBckdha. Full results of the $^{13}C$-MFA model.

| **3T3-L1 sgBCKDHa** | **Flux** | **95% Confidence Interval** | |
|---|---|---|---|
| **Pathway/Reaction** | **(nanomoles/well/hr)** | **Lower Bound** | **Upper Bound** |
| **Glycolysis** | | | |
| Glc.x -> G6P | 146.44 | 142.50 | 150.42 |
| G6P <-> F6P | 145.96 | 142.02 | 149.94 |
| G6P <-> F6P | 338.25 | 0.00 | Inf |
| F6P -> DHAP + GAP | 146.28 | 142.34 | 150.26 |
| DHAP <-> GAP | 146.28 | 142.34 | 150.26 |
| DHAP <-> GAP | 0.00 | 0.00 | Inf |
| GAP <-> 3PG | 292.73 | 284.84 | 300.67 |
| GAP <-> 3PG | 655.91 | 0.00 | Inf |
| 3PG -> PEP | 292.73 | 284.84 | 300.67 |
| PEP -> Pyr.c | 292.73 | 284.84 | 300.80 |
| Pyr.c <-> Lac | 139.59 | 129.18 | 149.83 |
| Pyr.c <-> Lac | 502.56 | 0.00 | Inf |
| Lac -> Lac.x | 139.59 | 129.18 | 149.83 |
| Pyr.c -> Ala | 6.54 | 0.76 | 7.64 |
| Pyr.m -> Ala | 0.00 | 0.00 | 5.73 |
| Ala -> Ala.x | 6.54 | 5.43 | 7.65 |
| **Pentose Phosphate Pathway** | | | |
| G6P -> P5P + CO2 | 0.48 | 0.42 | 0.48 |
| P5P + P5P <-> S7P + GAP | 0.16 | 0.14 | 0.16 |
| P5P + P5P <-> S7P + GAP | 0.00 | 0.00 | NaN |
| S7P + GAP <-> F6P + E4P | 0.16 | 0.14 | 0.16 |
| S7P + GAP <-> F6P + E4P | 0.00 | 0.00 | NaN |
| P5P + E4P <-> F6P + GAP | 0.16 | 0.14 | 0.16 |
| P5P + E4P <-> F6P + GAP | 0.00 | 0.00 | Inf |
| **Anaplerotic Reactions** | | | |
| Pyr.c -> Pyr.m | 146.60 | 134.52 | 159.13 |
| Pyr.m + CO2 -> Oac.m | 84.28 | 73.05 | 95.51 |
| Oac.c -> PEP + CO2 | 0.00 | 0.00 | 5.18 |
| Mal.m -> Pyr.m + CO2 | 98.22 | 88.30 | 110.01 |
| Mal.c -> Pyr.c + CO2 | 0.00 | 0.00 | 4.62 |
| PropCoA + CO2 -> SucCoA | 1.33 | 0.46 | 2.27 |
| Gln.c <-> Glu.c | 6.29 | -2.72 | 6.29 |
| Gln.c <-> Glu.c | 0.00 | 0.00 | 9.01 |
| Glu.c -> Glu.x | 0.18 | 0.17 | 0.19 |
| Glu.c <-> Glu.m | -18.00 | -28.50 | -11.68 |

| | | | |
|---|---|---|---|
| Glu.c <-> Glu.m | 0.00 | 0.00 | 9.01 |
| Glu.m <-> Akg.m | -11.51 | -21.98 | -5.22 |
| Glu.m <-> Akg.m | 42.62 | 22.21 | 74.63 |
| Akg.c <-> Glu.c | -24.11 | -34.61 | -17.79 |
| Akg.c <-> Glu.c | 11000.00 | 0.00 | Inf |
| Akg.m <-> Akg.c | 0.00 | -9.01 | 2.36 |
| Akg.m <-> Akg.c | 0.00 | 0.00 | 9.01 |
| Akg.c + CO2 <-> Cit.c | 24.11 | 17.80 | 30.85 |
| Akg.c + CO2 <-> Cit.c | 0.00 | 0.00 | 1.47 |
| Glu.m <-> Gln.m | -6.49 | -15.52 | -5.77 |
| Glu.m <-> Gln.m | 34.95 | 27.18 | 42.57 |
| Gln.x -> Gln.e | 12.78 | 12.06 | 13.50 |
| Gln.e -> Gln.c | 42.74 | 41.97 | 43.52 |
| Gln.c -> Gln.e | 29.96 | 29.58 | 30.35 |
| Gln.m <-> Gln.c | -6.49 | -15.52 | -5.77 |
| Gln.m <-> Gln.c | 72000.00 | 27.18 | Inf |
| **TCA Cycle** | | | |
| Pyr.m -> AcCoA.m + CO2 | 160.53 | 147.75 | 173.78 |
| AcCoAOther -> AcCoA.m | 19.29 | 15.58 | 23.24 |
| AcCoA.m + Oac.m -> Cit.m | 186.06 | 170.15 | 202.61 |
| Cit.m <-> Akg.m + CO2 | 55.55 | 48.94 | 62.80 |
| Cit.m <-> Akg.m + CO2 | 0.00 | 0.00 | 2.38 |
| Akg.m -> SucCoA + CO2 | 44.04 | 39.60 | 48.97 |
| SucCoA -> Suc | 45.37 | 40.55 | 50.72 |
| Suc <-> Fum.m | 45.37 | 40.55 | 50.72 |
| Suc <-> Fum.m | 0.01 | 0.00 | Inf |
| Fum.m <-> Mal.m | 45.37 | 40.55 | 50.72 |
| Fum.m <-> Mal.m | 83.23 | 0.00 | Inf |
| Mal.m <-> Oac.m | 101.78 | -30700.00 | 1080.00 |
| Mal.m <-> Oac.m | 1440.00 | 0.00 | Inf |
| Oac.m <-> Asp.m | 0.00 | -31300.00 | 940.84 |
| Oac.m <-> Asp.m | 0.00 | 0.00 | Inf |
| Mal.c <-> Oac.c | 0.00 | -31600.00 | Inf |
| Mal.c <-> Oac.c | 0.00 | 0.00 | Inf |
| Oac.c <-> Asp.c | 154.62 | -31200.00 | Inf |
| Oac.c <-> Asp.c | 51.29 | 0.00 | Inf |
| Asp.c -> Fum.c | 154.62 | 0.00 | Inf |
| Mal.c <-> Fum.c | -154.62 | -Inf | 0.00 |
| Mal.c <-> Fum.c | 4790.00 | 0.00 | NaN |
| Mal.c <-> Mal.m | 154.62 | -31100.00 | 30500.00 |

| | | | |
|---|---|---|---|
| Mal.c <-> Mal.m | 811.72 | 0.00 | Inf |
| Asp.m <-> Asp.c | 0.00 | -31300.00 | 940.84 |
| Asp.m <-> Asp.c | 0.00 | 0.00 | Inf |
| **BCAA Catabolism** | | | |
| Leu.x -> Leu | 5.63 | 5.15 | 6.11 |
| Leu <-> KIC | 2.08 | 1.54 | 2.65 |
| Leu <-> KIC | 2.33 | 0.00 | Inf |
| KIC <-> IsoVCoA + CO2 | 2.08 | 1.54 | 2.65 |
| KIC <-> IsoVCoA + CO2 | 0.05 | 0.00 | 0.23 |
| IsoVCoA + CO2 <-> HMGCoA | 2.08 | 1.54 | 2.65 |
| IsoVCoA + CO2 <-> HMGCoA | 1.34 | 0.00 | 920000.00 |
| HMGCoA -> AcCoA.m + AcCoA.m + AcCoA.m | 2.08 | 1.54 | 2.65 |
| LeuT -> Leu | 1.37 | 1.24 | 1.52 |
| Leu -> LeuP | 4.93 | 4.10 | 5.73 |
| Val.x -> Val | 3.33 | 2.86 | 3.79 |
| ValT -> Val | 0.69 | 0.58 | 0.81 |
| Val -> ValP | 0.00 | 0.00 | 3.17 |
| Val <-> IsoBCoA + CO2 | 4.02 | 2.67 | 4.58 |
| Val <-> IsoBCoA + CO2 | 0.30 | 0.21 | 0.39 |
| IsoBCoA <-> MMA | 4.01 | 1.00 | 4.58 |
| IsoBCoA <-> MMA | 0.00 | 0.00 | Inf |
| MMA -> PropCoA + CO2 | 4.01 | 1.00 | 4.58 |
| **Fatty Acid Synthesis** | | | |
| Cit.m -> Cit.c | 130.51 | 115.67 | 145.95 |
| Cit.c -> AcCoA.c + Oac.c | 154.62 | 140.14 | 169.69 |
| AcCoA.c + AcCoA.c + AcCoA.c + AcCoA.c + AcCoA.c + AcCoA.c + AcCoA.c + AcCoA.c -> Palm.s | 16.85 | 14.52 | 20.99 |
| Palm.s -> Palm | 16.85 | 14.52 | 20.99 |
| Palm.d -> Palm | 7.31 | 6.24 | 8.97 |
| PropCoA + AcCoA.c + AcCoA.c + AcCoA.c + AcCoA.c + AcCoA.c + AcCoA.c -> C15FA.s | 2.68 | 0.00 | 3.73 |
| C15FA.s -> C15FA | 2.68 | 0.00 | 3.73 |
| C15FA.d -> C15FA | 1.04 | 0.00 | 1.64 |
| C15FA + AcCoA.c -> C17FA.s | 3.73 | 1.20 | 5.25 |
| C17FA.s -> C17FA | 3.73 | 1.20 | 5.25 |
| C17FA.d -> C17FA | 4.11 | 1.03 | 6.00 |

| | | | |
|---|---|---|---|
| IsoBCoA + AcCoA.c + AcCoA.c + AcCoA.c + AcCoA.c + AcCoA.c + AcCoA.c + AcCoA.c + AcCoA.c -> IsoC16.s | 0.00 | 0.00 | 3.17 |
| IsoC16.s -> IsoC16 | 0.00 | 0.00 | 3.17 |
| IsoC16.d -> IsoC16 | 0.00 | 0.00 | Inf |
| **Dilution/Mixing** | | | |
| 0*Pyr.c -> Pyr.mnt | 0.00 | 0.00 | 0.02 |
| 0*Pyr.m -> Pyr.mnt | 1.00 | 0.98 | 1.00 |
| 0*Mal.c -> Mal.mnt | 0.00 | 0.00 | 1.00 |
| 0*Mal.m -> Mal.mnt | 1.00 | 0.00 | 1.00 |
| 0*Asp.c -> Asp.mnt | 0.90 | 0.00 | 1.00 |
| 0*Asp.m -> Asp.mnt | 0.10 | 0.00 | 1.00 |
| 0*Fum.m -> Fum.mnt | 0.03 | 0.00 | 1.00 |
| 0*Fum.c -> Fum.mnt | 0.97 | 0.00 | 1.00 |
| 0*Cit.m -> Cit.mnt | 0.57 | 0.33 | 0.75 |
| 0*Cit.c -> Cit.mnt | 0.43 | 0.25 | 0.67 |
| 0*Glu.m -> Glu.mnt | 1.00 | 0.67 | 1.00 |
| 0*Glu.c -> Glu.mnt | 0.00 | 0.00 | 0.33 |
| 0*Gln.m -> Gln.mnt | 0.00 | 0.00 | 1.00 |
| 0*Gln.c -> Gln.mnt | 1.00 | 0.00 | 1.00 |
| 0*Akg.m -> Akg.mnt | 0.63 | 0.48 | 0.81 |
| 0*Akg.c -> Akg.mnt | 0.37 | 0.19 | 0.52 |
| Pyr.mnt -> Pyr.fix | 1.00 | 1.00 | 1.00 |
| Asp.mnt -> Asp.fix | 1.00 | 1.00 | 1.00 |
| Mal.mnt -> Mal.fix | 1.00 | 1.00 | 1.00 |
| Fum.mnt -> Fum.fix | 1.00 | 1.00 | 1.00 |
| Cit.mnt -> Cit.fix | 1.00 | 1.00 | 1.00 |
| Akg.mnt -> Akg.fix | 1.00 | 1.00 | 1.00 |
| Glu.mnt -> Glu.fix | 1.00 | 1.00 | 1.00 |
| Gln.mnt -> Gln.fix | 1.00 | 1.00 | 1.00 |
| SSR | 617 | 580 | 721 |

**Supplementary Table 4. CRISPR/Cas9 target sequences.** Sense and anti-sense single-guide sequences of Control, Bckdha, and Acad8 guides used in pooled cell culture experiments

| Guide Target | Sense | Anti-sense |
|---|---|---|
| Control guide | CACCGGCCGTGTTGCTGGATACGCC | AAACGGCGTATCCAGCAACACGGCC |
| Bckdha guide 1 | CACCGCAGCGAAATTGAAACCGGCG | AAACCGCCGGTTTCAATTTCGCTGC |
| Bckdha guide 2 | CACCGTGAGGGATCTGCGTGGCCAG | AAACCTGGCCACGCAGATCCCTCAC |
| Bckdha guide 3 | CACCGCATGACCAACTATGGCGAGG | AAACCCTCGCCATAGTTGGTCATGC |
| Acad8 guide 1 | CACCGCCTTCCGCATCACATCCACA | AAACTGTGGATGTGATGCGGAAGGC |
| Acad8 guide 2 | CACCGGCCAACAGGATTGGGACCG | AAACCGGTCCCAATCCTGTTGGCC |
| Acad8 guide 3 | CACCGAGGTGAGTCAGACATCTATG | AAACCATAGATGTCTGACTCACCTC |

**Supplementary Table 5. Primer sequences.** Forward and reverse sequences of primers used to quantify expression of genes using quantitative RT-qPCR.

| Gene name | Forward Sequence | Reverse Sequence |
|---|---|---|
| 18S rRNA | AGTCCCTGCCCTTTGTACACA | CGATCCGAGGGCCTCACTA |
| Adiponectin (*AdipoQ*) | GACACCAAAAGGGCTCAGGA | GCCCTTCAGCTCCTGTCATT |
| Carbohydrate response element binding protein (*Chrebp*) | CACTCAGGGAATACACGCCTAC | ATCTTGGTCTTAGGGTCTTCA |
| Fatty acid binding protein 4 (*Fabp4*) | AGAAGTGGGAGTGGGCTTTG | CCAGCTTGTCACCATCTCGT |
| Glucose transporter type 4 (*Glut4*) | AGCCTCTGATCATCGCAGTG | ACTAAGAGCACCGAGACCAAC |
| Peroxisome Proliferator-Activated Receptor Gamma Coactivator 1-Alpha (Pgc1a) | CCCTGCCATTGTTAAGACC | TGCTGCTGTTCCTGTTTTC |
| Peroxisome proliferator-activated receptor γ (*PPARg*) | TTCGCTGATGCACTGCCTAT | ACAGACTCGGCACTCAATGG |
